# The complex genomic basis of rapid convergent adaptation to pesticides across continents in a fungal plant pathogen

**DOI:** 10.1101/2020.07.24.220004

**Authors:** Fanny E. Hartmann, Tiziana Vonlanthen, Nikhil Kumar Singh, Megan McDonald, Andrew Milgate, Daniel Croll

## Abstract

Convergent evolution leads to identical phenotypic traits in different species or populations. Convergence can be driven by standing variation allowing selection to favor identical alleles in parallel or the same mutations can arise independently. However, the molecular basis of such convergent adaptation remains often poorly resolved. Pesticide resistance in agricultural ecosystems is a hallmark of convergence in phenotypic traits. Here, we analyze the major fungal pathogen *Zymoseptoria tritici* causing serious losses on wheat and with parallel fungicide resistance emergence across continents. We sampled three population pairs each from a different continent spanning periods early and late in the application of fungicides. To identify causal loci for resistance, we combined knowledge from molecular genetics work and performed genome-wide association studies (GWAS) on a global set of isolates. We discovered yet unknown factors in azole resistance including membrane stability functions. We found strong support for the ‘hotspot’ model of resistance evolution with parallel changes in a small set of loci but additional loci showed more population-specific allele frequency changes. Genome-wide scans of selection showed that half of all known resistance loci were overlapping a selective sweep region. Hence, the application of fungicides was one of the major selective agents acting on the pathogen over the past decades. Furthermore, loci identified through GWAS showed the highest overlap with selective sweep regions underlining the importance to map phenotypic trait variation in evolving populations. Our population genomic analyses showed that both *de novo* mutations and gene flow likely contributed to the parallel emergence of resistance.

## Introduction

A major interest in evolutionary biology is how the same phenotype can arise in distinct populations or lineages (Losos, 2011). Repeated adaptive evolution can produce impressive reproducibility in the emergence of the same trait. Previously, the terms convergent and parallel adaptive evolution were used to distinguish the evolution of the same trait in distant taxa (often based on distinct mechanisms) from the evolution of the same trait among closely related species. This distinction has only weak support though and we use here the term convergent for all cases of repeated evolution of the same trait (Arendt & Reznick, 2008). Historically, investigations of convergent evolution largely focused on long-separated species. However, the advent of population genomics and experimental approaches produced a series of well-supported cases of convergent evolution among closely related species or even populations of the same species (Bolnick, Barrett, Oke, Rennison, & Stuart, 2018). Convergent evolution among populations can proceed either through the independent emergence of mutations in either the same or different genes (Chan et al., 2010; Pearce et al., 2009; Stern, 2013; Tishkoff et al., 2007). Alternatively, convergent evolution can arise from standing variation, hence the mutations were either present in an ancestor or were spread among populations through gene flow (Colosimo, 2005; Heliconius Genome Consortium, 2012; Roesti, Gavrilets, Hendry, Salzburger, & Berner, 2014; Song et al., 2011). Elucidating the genetic basis of convergent evolution can provide fundamental insights into the role of gene flow, pleiotropy and mutation limitation in adaptive evolution (Orr & Allen Orr, 2005). However, few experimental systems allow both the tracking of adaptive mutations underlying phenotypic trait convergence and population genetic analyses of such mutations limiting our understanding of rapid convergent evolution.

The emergence of pesticide resistance in agricultural ecosystems is a hallmark system for investigating convergent evolution at the molecular and population genetic level (Baucom, 2019; Hawkins, Bass, Dixon, & Neve, 2019). In weeds evolving resistance to herbicides, often the same genetic changes are observed across lineages of the same species (Menchari et al., 2006; Powles & Yu, 2010). The underlying mutations are mostly focused on the gene encoding a key enzyme being perturbed by a given herbicide. Similarly, convergent evolution in fungal pathogens has led to repeated changes in the gene encoding a key sterol biosynthesis enzyme targeted by azole fungicides (Lucas, Hawkins, & Fraaije, 2015). The highly reproducible involvement of the same pathways contributing to pesticide resistance in plants and fungi, respectively, supports a “hotspot” model of resistance evolution (Martin & Orgogozo, 2013). Furthermore, the “hotspot” genes often show strong mutational constraints driven by pleiotropic effects due to the essential functions of the encoded protein. Key examples of hotspots of adaptation include the acetolactate synthase (ALS) gene which encodes an enzyme catalyzing the first step in the biosynthesis of branched amino acids targeted by multiple classes of herbicides (Shimizu et al., 2008). In fungi, azole fungicides act on the sterol biosynthesis cytochrome P450 monooxygenase CYP51 (Cools, Hawkins, & Fraaije, 2013). In both weeds and fungal pathogens, nearly identical mutations have been observed across species following gains of resistance (Cools et al., 2013; Délye, Jasieniuk, & Le Corre, 2013; Mohd-Assaad, McDonald, & Croll, 2016; Parker et al., 2014). The high degree of convergence at the level of individual mutations suggests that pleiotropy plays a key role in shaping the spectrum of observed resistance mutations. Analysis of large numbers of natural mutants of the wheat pathogen *Zymoseptoria tritici* revealed that the mutational pathway to resistance is indeed highly constrained and mutations largely follow a predictable sequence of appearance (Hawkins et al., 2019). Similarly, mutations in the ALS gene are strongly biased towards two specific amino acid changes (Tranel, Wright, & Heap, 2020). Beyond convergent evolution in hotspot genes, plants and fungi gained resistance by a variety of additional mechanisms including detoxification, overexpression of the target gene and re-wiring of metabolic pathways (Cools, Bayon, Atkins, Lucas, & Fraaije, 2012; Molin, Yaguchi, Blenner, & Saski, 2020). Compared to target-site resistance, such non-target site resistance can lead to convergence at the phenotypic level but is less likely to lead to convergence at the molecular level (Baucom, 2019; Losos, 2011). Hence, disentangling the relative contributions of different resistance mechanisms is crucial for understanding convergent evolution against pesticides.

Resistance to fungicides by fungal pathogens is a major threat to agricultural yields but also human health (Fisher et al., 2012). In both clinical and agricultural practice, azoles are the most commonly used class of fungicides (*i.e.* imidazoles and triazoles) (Azevedo, Faria-Ramos, Cruz, Pina-Vaz, & Rodrigues, 2015; Cools et al., 2013). To protect crops, strobilurins, succinate dehydrogenase inhibitors (SDHI) and benzimidazoles (MBC) are commonly used in addition to or in combination with azoles. Fungicide resistance breakdown has been observed in many plant pathogens with the most repeatable breakdown being the gain of resistance to strobilurin (QoI) fungicides (Hawkins et al., 2019). A single point mutation in the mitochondrial gene *cytb* confers resistance and the same mutation arose independently across species or even populations, for example populations of the globally distributed pathogen *Z. tritici* (Torriani, Brunner, McDonald, & Sierotzki, 2009). This pathogen is the causal agent of Septoria tritici blotch disease on wheat, causing dramatic annual economic losses (Torriani et al., 2015). The wheat pathogen has a well documented history of fungicide resistance that spans the globe. For some fungicides, resistance emerged very rapidly (*i.e.* QoI), whereas for other modes of action the resistance has arisen more gradually (*i.e.* azoles) (Torriani et al., 2015). The pathogen originated in the Middle East and spread first to Europe later followed by colonization events of the Americas and Oceania (Zhan, Pettway, & McDonald, 2003). Strobilurin and azole resistance emerged across continents often following only a few years after the first fungicide applications (Estep et al., 2015; Torriani et al., 2009). It is estimated that ~50% of all fungicides sprayed on cereals in Europe target the pathogen largely due to the loss in efficacy of most fungicides. Resistance to SDHI was recently discovered in European populations (Rehfus, Strobel, Bryson, & Stammler, 2018). Azole resistance emerged in Australia independently multiple times with possible contributions from gene flow from European populations (McDonald et al., 2019). The pathogen harbors large, panmictic populations (Singh, Chanclud, & Croll, 2020; Zhan et al., 2003) with the ability to rapidly surmount host resistance by fixing mutations in key virulence genes (Cowger, Hoffer, & Mundt, 2000; Hartmann, Sánchez-Vallet, McDonald, & Croll, 2017; Meile et al., 2018; Zhong et al., 2017). *Z. tritici* received also considerable attention to dissect the molecular genetic basis of fungicide resistance with multiple large-scale mutant screens (Cools & Fraaije, 2013). *Z. tritici* shows a broad range of complementary resistance mechanisms including target site resistance supporting the “hotspot” model and an array of detoxification mechanisms such as the upregulation of transporter genes (Omrane et al., 2017, 2015).

In this study, we retraced the emergence of fungicide resistance on three continents by performing genome sequencing and association mapping analyses of a total of 356 isolates. We first aimed to comprehensively catalogue loci underlying resistance by combining previous knowledge from molecular genetics work with a genome-wide association study (GWAS) of a global set of isolates. Second, we aimed to quantify the parallelism in adaptive allele frequency changes across populations to assess how likely populations gained resistance independently. Finally, we tested whether recent selection for fungicide resistance showed convergent or distinct genomic signatures across population pairs.

## Materials and Methods

### Fungal isolate collection and genomic data

We analyzed a total of 356 *Z. tritici* isolates sampled in three main regions including Australia, Switzerland and the United States (US; Oregon) (Supplementary Table S1). In each region, two samplings were made at two different time points corresponding to absent or low degrees of fungicide application versus higher levels of fungicide applications. In Australia, the older collection was made in 2001 in Wagga Wagga (Southeastern Australia) and the more recent collection was made in 2014 in Tasmania (McDonald et al., 2019; Zhan et al., 2003). The older Swiss collection was made in 1999 near Winterthur (cantonZurich) at a time when azoles were in use already and the more recent collection was made in 2016 in nearby Eschikon (both canton Zurich) in a field with extensive application of fungicides (Oggenfuss, Badet, Wicker, & Hartmann, 2020; Zhan et al., 2003). The older collection from the United States was made in 1990 in Willamette Valley (Oregon) before the use of azoles in the region and the more recent collection was made in 2015 in the same location following a decade of increasing fungicide applications (Estep et al., 2015).

### DNA extraction, genome sequencing and variant calling

We used *ca.* 100 mg of lyophilized spores to extract high-quality genomic DNA following the Qiagen DNAeasy Plant Mini Kit extraction protocol. Genomic DNA was sequenced on a Illumina HiSeq 4000 using a 100 bp paired-end cycle protocol. All raw reads are deposited on the NCBI Short Read Archive under the BioProject PRJNA596434 (Oggenfuss et al., 2020; Singh et al., 2020), PRJNA327615 (Hartmann, McDonald, & Croll, 2018; Hartmann et al., 2017) and PRJNA480739 (McDonald et al., 2019). Raw sequencing reads were quality-trimmed (Illuminaclip = TruSeq3-PE.fa:2:30:10, leading = 10, trailing = 10, sliding window = 5:10, minlen = 50) using Trimmomatic v0.32 (Bolger, Lohse, & Usadel, 2014) Trimmed reads were aligned to the fully assembled reference genome of the species (isolate IPO323) (Goodwin et al., 2011) using bowtie2 v. 2.4.1 (Langmead & Salzberg, 2012). We called single-nucleotide polymorphisms (SNPs) using the Genome Analysis Toolkit v. 4.0.1.2 (McKenna et al., 2010) running first HaplotypeCaller (ploidy = 1) on each individual and then generating combined variant calls using GenotypeGVCF. Variant calls were hard filtered removing any variant satisfying any of the following conditions: QD < 5.0; QUAL < 1000.0; MQ < 20.0; −2 > ReadPosRankSum > 2.0; −2 > MQRankSum > 2.0; −2 > BaseQRankSum > 2.0. We identified the most likely ancestral state at SNP loci using whole-genome sequencing data of the two closest known sister species of *Z. tritici* : four *Z. pseudotritici* isolates (STIR04_2.2.1, STIR04_3.11.1, STIR04_5.3 and STIR04_5.9.1) and four *Z. ardabiliae* isolates (STIR04_1.1.1, STIR04_1.1.2, STIR04_3.13.1 and STIR04_3.3.2) (Stukenbrock et al., 2011, 2010). In addition to the filters above, we retained the ancestral state at SNPs if the SNP genotyping rate was >50% among sister species and no polymorphism within and between sister species of *Z. tritici* was found.

### Population structure analyses

We analyzed population structure using SNPs called on all chromosomes (excluding the mitochondrial genome). We retained only bi-allelic SNPs with a genotyping rate >80%. We used two complementary approaches. First, we performed a principal component analysis (PCA) using the --pca command of the software PLINK v1.9 (Purcell et al., 2007). Then, we used the model-based clustering approach implemented in FastStructure v1.0 (Raj, Stephens, & Pritchard, 2014). For this, we used only 2,364 genome-wide at a distance of 15 kb along the chromosomes constituting a set of most likely unlinked SNPs. Previous estimates of linkage disequilibrium decay in *Z. tritici* populations revealed that linkage disequilibrium decays at least at a distance about a magnitude smaller (Hartmann et al., 2018, 2017). We used the R package {Pophelper} v1.2.0 (https://github.com/royfrancis/pophelper) to generate barplots. Cluster assignment probabilities were computed using the CLUMPP program (Jakobsson & Rosenberg, 2007) implemented in the R package {Pophelper}. We used the “chooseK.py” script of FastStructure and visual inspection to assess the most relevant number of groups (K).

### Fungicide resistance assay

A microtiter plate assay was performed to quantify azole resistance based on propiconazole. All populations were screened with the exception of the collection from Tasmania (Australia) and the recent Swiss population using an established protocol (Cools et al., 2011). Growth inhibition was tested over 12 different concentrations of propiconazole (Syngenta Inc., Stein, Switzerland) with concentrations including 1.5, 0.55, 0.20, 0.072, 0.042, 0.025, 0.015, 0.0086, 0.0051, 0.00017, 0.00006 mg liter^−1^ and a control without fungicide. We placed 100 μl of Sabouraud-dextrose liquid medium (SDLM; Oxoid, Basingstoke, England) amended with the respective concentration of propiconazole into microtiter wells. Wells were supplemented with 100 μl of spore suspensions at a concentration of 2.5 × 10^4^ spores per ml. Microtiter plates were shaken at low speed for one minute, sealed with parafilm and incubated in the dark for four days at 21°C and 80% relative humidity. Fungal growth was measured with an Elisa plate reader (MR5000, Dynatech) by assessing the optical density at 605 nm. The growth inhibition at different propiconazole concentrations was used to estimate the dose-response curves of different isolates. The half-maximal concentrations (EC_50_) were estimated based on a 4-parameter logistic curve using the R package {drc} (Ritz and Streibig, 2005)

### Phenotype-genotype associations for fungicide resistance and confirmed resistance mutations

We identified phenotype-genotype associations based on a mixed linear model in TASSEL v 5 using a kinship matrix as a random factor to correct for population structure (Bradbury et al., 2007). We performed two separate GWAS: on a total of 134 isolates belonging to the two populations from the United States and on a total of 211 isolates including all populations except the Australian collection from Tasmania and the more recent collection from Switzerland. We retained significantly associated loci based on the Bonferroni threshold α = 0.05 and the false discovery rate (FDR) threshold of 5%. FDR calculations were done using the R package {qvalue} v2.6.0 (Storey, Bass, Dabney, & Robinson, 2015). In addition to loci retrieved from GWAS, we combined information for previously reported and functionally confirmed resistance mutations to strobilurin (QoI), benzimidazole (MBC), azoles and succinate dehydrogenase inhibitors (SDHI). The genomic location of mutations and relevant publications are shown in Supplementary Table S2.

### Genetic diversity and selection analyses

We computed linkage disequilibrium heatmaps and decay within each population using the option --hap-r2 of the vcftools v0.1.15 program (Danecek et al, 2011). We selected SNPs without missing data and filtered for a minor allele frequency >0.05. For decay analyses, we analyzed the largest chromosome (chromosome 1) only. We computed allele frequency and allele counts within populations using the -freq and -count options of vcftools. We computed per-gene diversity statistics such as nucleotide diversity per site (π), Tajima’s *D* within populations, relative divergence index (*F*_*ST*_) and the absolute divergence index (*d*_*XY*_) between populations using the R package PopGenome v2.7.5 (Pfeifer, Wittelsbürger, Ramos-Onsins, & Lercher, 2014). To identify signatures of recent positive selection, we used two extended haplotype homozygosity (EHH) based-tests (Sabeti et al., 2007) implemented in the R package REHH v2.0 (Gautier, Klassmann, & Vitalis, 2017). First, we analyzed signatures of selective sweeps in each of the three more recent populations by computing the integrated haplotype score (iHS) statistic. The iHS statistic aims to detect abnormally long haplotype blocks by comparing the integrated EHH of the ancestral allele and the integrated EHH of the derived allele at each SNP. We used a maximal distance gap of 20 kb for the Swiss and the United States populations and 500 kb for the Australian population as estimates of linkage disequilibrium decay differed between geographical regions. Then, we performed cross-population extended haplotype homozygosity (XP-EHH) scans between the older and more recent population in each geographical region. The XP-EHH test compares the profile of EHH between pairs of clusters at each focal SNP to detect regions under positive selection in one population compared to another population. We used as an outlier detection threshold the 99.9^th^ percentile of the distribution of absolute iHS and XP-EHH values. Therefore, we performed six independent genome-wide scans in total: three within-population scans and three inter-populations scans. To define the size of selective sweep regions to consider, we used information on linkage disequilibrium decay. In the Swiss and the United States populations, we clustered significant SNPs in separate sweep regions if the distance between significant SNPs was >5 kb and added 2.5 kb around the significant SNPs to define the region. To define selective sweep regions in the Australian populations, we clustered significant SNPs in separate sweep regions if the distance between significant SNPs was >50 kb and added 25 kb +/− around the SNPs.

## Results

### Parallel emergence of fungicide resistance in population pairs

We analyzed populations of the wheat pathogen *Z. tritici* in three distinct geographic regions including North America (Oregon, United States), Central Europe (Switzerland) and Australia having experienced an increase in fungicide applications over the last three decades (Estep et al., 2015; McDonald et al., 2019; Torriani et al., 2015) (Supplementary Table S1; Fig 1A, Supplementary Fig S1). The North American and Central European populations were sampled at the same or nearby site at an interval of 25 and 17 years, respectively. The older Australian population was sampled in 2001 in South Eastern Australia and the more recent population was sampled in 2014 in Tasmania further south. We quantified the shift in fungicide resistance in the pair of North American populations collected in the United States. The population showed a strong shift from sensitivity to resistance to the commonly sprayed azole propiconazole (Fig 1B).

**Figure 1:**
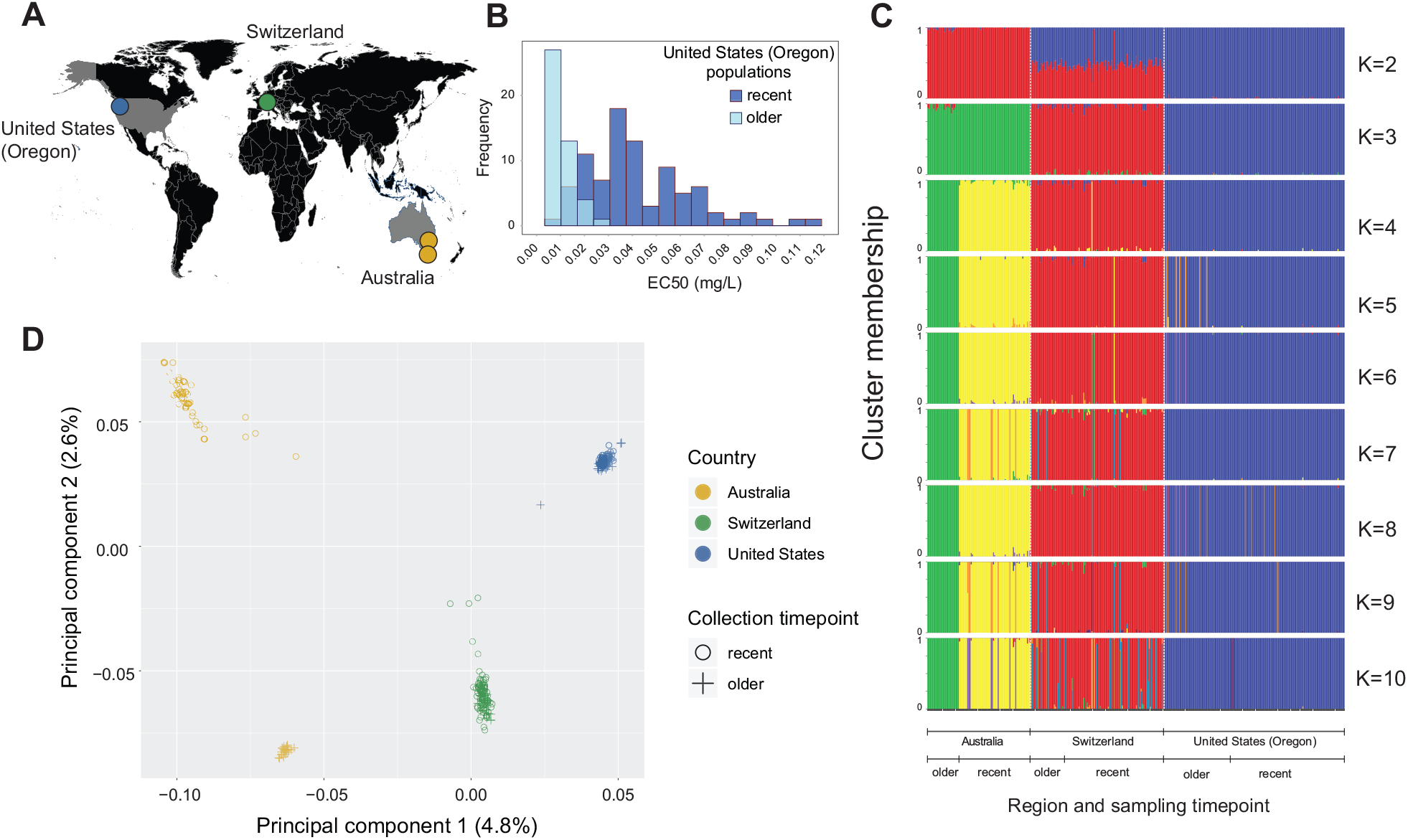
Population analyses of 356 genomes *of Zymoseptoria tritici.* A) Sampling locations in three geographical regions and at two different time points. B) Fungicide resistance to propiconazole of the historical and recent populations sampled in the United States (Oregon) expressed as the concentration of the half-maximal growth (EC50). C) Results of FastStructure for K ranging from 2-10 based on 2,364 genome-wide, unlinked SNPs. Genotypes were grouped according to their geographic origin and sampling time point. D) Principal component analysis on 1,380,931 genome-wide SNPs. The first and second axis of the principal components are shown and the percentage of variance explained by each axis is indicated in brackets.

The worldwide collection included 356 isolates, which were all whole-genome sequenced using Illumina. A principal component analysis (PCA) and unsupervised clustering using FastStructure revealed a clear geographic separation of the different genotypes (Fig 1C-D). Australia, Switzerland, and the United States are clearly distinguished by the first two axes of the PCA and were assigned to distinct clusters at K=3 (Fig 1C-D). We found no population substructure between the older and more recent populations sampled in Switzerland and the United States (Fig S2 B-C). In contrast, the populations sampled in Australia were clearly distinct both when analyzed jointly with the other populations and alone (Fig 1 C-D; Supplementary Fig S2). The genetic differentiation among the Australian isolates could be largely explained by the geographically distinct sampling and but also possible gene flow from Europe into the Tasmanian population. In the FastStructure analysis, K=4 provided the lowest model complexity while maximizing the marginal likelihood. We considered therefore four population genetic units: all isolates from Switzerland, all isolates from the United States and separately the older and more recent isolates sampled in Australia.

Consistent with the analyses of population structure, linkage disequilibrium decay occurs at short distances in the Switzerland and the United States populations. The *r*^2^ reaches 0.2 at ~600 bp and ~400 bp in the older and more recent Swiss populations, respectively (Supplementary Fig S3). The *r*^2^ reaches 0.2 at ~1.8 kb and ~1 kb in the older and more recent United States populations, respectively. The older Australian population has a substantially slower decay in linkage disequilibrium (*r*^2^ reaching 0.2 at ~8.6 kb), which is likely due to the founder effect during the colonization of Australia (Hartmann et al., 2017; Zhan et al., 2003). The more recent Australian population from Tasmania shows an even slower decay with *r*^2^ reaching 0.2 at ~50 kb caused possibly by recent admixture from Europe as indicated by the population structure analyses (Fig. 1C-D). The pathogen recently evolved resistance to a commonly used azole fungicide across continents. The genetic structure suggests that at least in Europe and North America gene flow may have played only a minor role in bringing in resistance alleles because the overall genetic structure has remained stable. Resistance in Australia may have evolved jointly from standing variation and gene flow from Europe.

### Mapping the genetic basis of fungicide resistance

The parallel evolution of resistance raises the question whether the underlying mutations are congruent or distinct among continents requiring precise knowledge of causal loci (Fig 2A). Resistance to fungicides most often involves mutations in the genes underlying the proteins targeted by the chemical (Cools et al., 2013). However, additional loci can play an important role *e.g.* in cell detoxification processes or by creating redundancy in the targeted protein. To comprehensively investigate parallel resistance evolution, we combined knowledge of confirmed fungicide resistance loci with GWAS. *Z. tritici* evolved resistance to at least four main classes of fungicide (Cools & Fraaije, 2013). We analyzed polymorphism at loci with known mutations causing resistance to strobilurins (mitochondrial gene *cytb*), azoles (*CYP51*) and SDHIs (*sdh* genes 2-4) and carbendazim (beta-tubulin) (Supplementary Table S2). We excluded complex resistance mutations such as the insertion of transposable elements upstream of the gene encoding MgMFS1 (Omrane et al., 2017) because these would be potentially unreliable to score from genome sequencing data. To expand knowledge on the genetic architecture of emerging azole fungicide resistance, we performed GWAS on the pair of populations collected in North America (*n* = 134 isolates). The low degree of differentiation between the older and more recent collection reduces confounding effects of relatedness and resistance. We mapped 141 significantly associated SNPs (Bonferroni alpha = 0.05) and 60 additional SNPs at false discovery rate (FDR) of 5% (Supplementary Table S3A; Fig 2B). The significant SNPs clustered in 8 genomic regions on chromosomes 1, 3, 5, 7, 12 and 13. A total of 148 SNPs were matching a gene. Among the 53 intergenic SNPs, 29 SNPs were located at less than 2.5 kb of a gene. We expanded our GWAS to cover a previously established worldwide collection of *Z. tritici* isolates (*n* = 211) spanning all continents affected by the pathogen (Hartmann et al., 2017). We identified 134 significantly associated SNPs at a Bonferroni alpha = 0.05 threshold and 63 additional SNPs at a 5% FDR (Supplementary Table S3B; Supplementary Fig 4B). The GWAS on the worldwide collection confirmed the same regions as in the first GWAS with the exception of a region on chromosome 1 (at 5.795 Mb) and a region on chromosome 7 (at 1.371 Mb). Hence, the worldwide GWAS generated a consistent set of loci largely matching the narrower GWAS panel focused on the North American population pair.

**Figure 2 :**
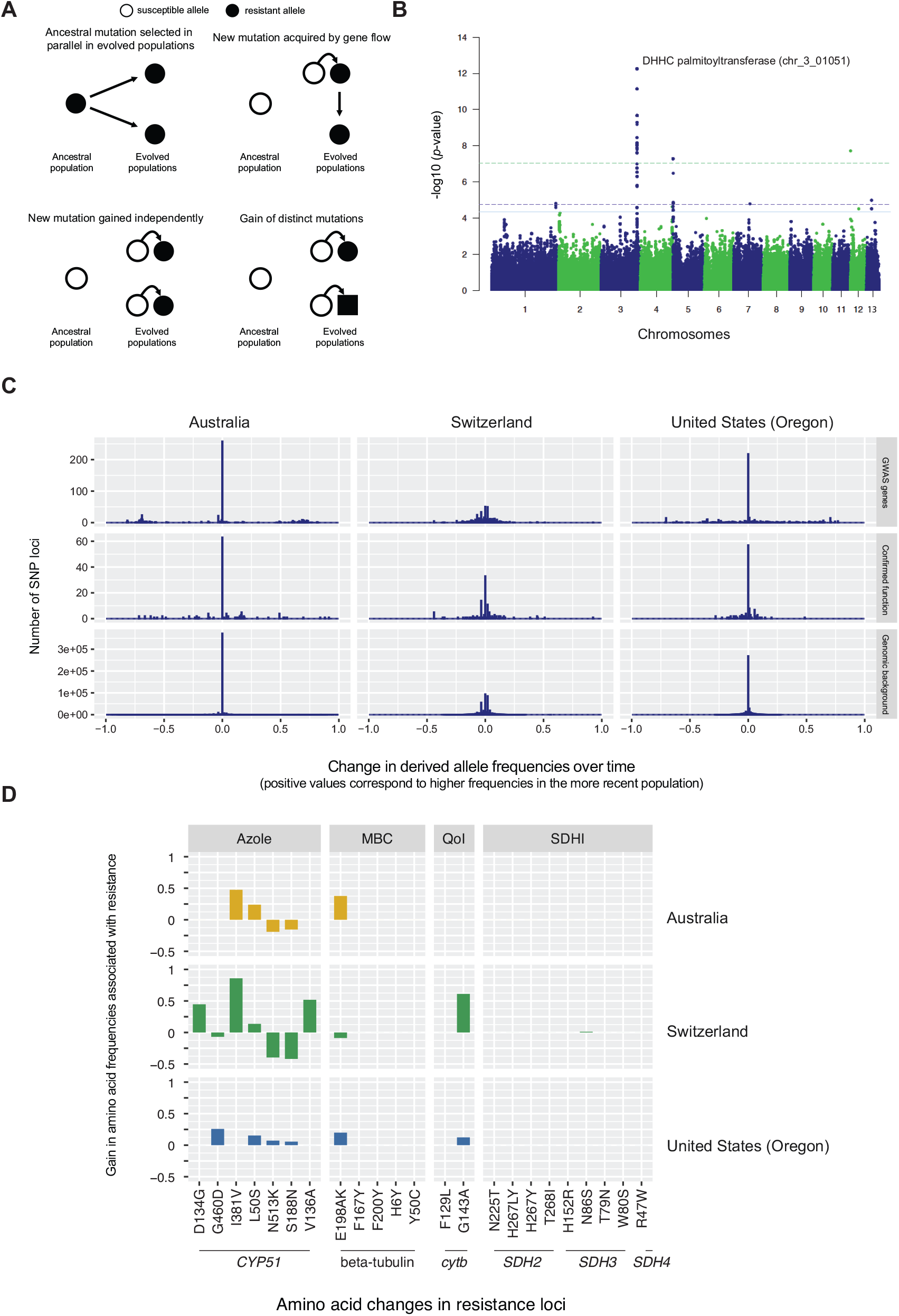
Association mapping of resistance emergence and parallel changes in resistant allele frequencies. A) Conceptual view of four possible origins of resistance mutations in evolved populations. Filled circles or square represent resistance mutations. B) Manhattan plot of the genome-wide association mapping analyses (GWAS) for the combined older and more recent populations from the United States (Oregon; *n* = 134). The Bonferroni (alpha = 0.05), FDR 5% and FDR 10% thresholds are shown with dotted green lines, dotted blue and solid blue lines, respectively. C) Allele frequency changes are expressed as the difference in derived allele frequencies between older and more recent populations. Positive values correspond to higher frequencies of the derived allele in more recent populations. SNPs in genes identified through GWAS and functionally confirmed genes are shown separately. The genomic background corresponds to all remaining genes. D) Changes in amino acid frequencies in populations between older and more recent populations. Positive frequency changes correspond to gains in the amino acid associated with higher resistance. Proteins CYP51, beta tubulin, cytb and SDH2-4 are targeted by azole, MBC, QoI and SDHI fungicides, respectively. Genotypes are shown in Supplementary Table S8.

The chromosomal region most significantly associated with azole fungicide resistance was identified by 20 adjacent SNPs on chromosome 3 in complete linkage disequilibrium (Supplementary Table S4). The locus matches the gene *Zt09_chr_3_01051*, which encodes a DHHC palmitoyl transferase. This enzyme is responsible for attaching palmitate to membrane proteins (Fukata, Fukata, Adesnik, Nicoll, & Bredt, 2004; Linder & Deschenes, 2007). The membrane-associated gene function may help to alleviate the deleterious effects of azoles on the sterol biosynthesis and membrane structure. Three nearby genes (*Zt09_chr_3_1048*, *1052* and *1053*) also harbored significantly associated SNPs. These associations most likely arose through linkage disequilibrium with the most significant SNP. Additional associations were found for SNPs in four genes including a SNP in the *CYP51* gene encoding the protein target of azole fungicides. We also identified a significant association with the gene *AvrStb6*, which encodes a major virulence factor (Zhong, Marcel, 2017). This association most likely arose because the pathogen is known to have concurrently evolved fungicide resistance as well as surmounted host resistance based on *AvrStb6* mutations in the United States (Cowger et al., 2000). We excluded the locus *AvrStb6* from further analyses. Hence, our search for mutations in experimentally confirmed resistance genes and previously unknown resistance genes mapped through GWAS produced a combined list of 13 non-redundant loci (Supplementary Table S2, S4).

### Adaptive allele frequency changes

Precise knowledge of resistance mutations enables tracking of adaptive allele frequency changes over time. We first focused globally on allele frequency changes at SNPs in genes associated with fungicide resistance (Fig 2C). To establish background levels of allele frequency changes, we analyzed genome-wide SNPs where we could ascertain the ancestral state (546'669 SNPs; see Methods for details). Genome-wide polymorphisms showed only very minor changes in allele frequencies for the Australian and the United States population pairs, and slightly more pronounced allele frequency shifts in the Swiss population pair (Fig 2C). In contrast, both fungicide resistance loci identified by GWAS and functionally confirmed loci showed stronger changes in allele frequencies in all population pairs (Fig 2C; Supplementary Figure 5A-B). The overall higher variance in allele frequencies at fungicide resistance loci compared to the genomic background suggests that these genes indeed contributed to resistance evolution and exceed effects of other evolutionary processes, such as genetic drift or population admixture. Interestingly, the changes in allele frequencies at resistance loci have been close to symmetrical in regards to changes in derived alleles. Even though functionally relevant alleles for resistance are expected to be derived, linkage disequilibrium with additional mutations at resistance loci likely underpins the overall symmetrical increases in both ancestral and derived alleles.

To investigate protein-level changes relevant for resistance, we analyzed amino acid frequencies among population pairs in the ‘hotspot’ gene for each fungicide class (Fig 2D; Supplementary Table S5; Supplementary Figure 5C). We used both information from molecular genetics studies and ancestral state reconstruction from sister species to define the most likely susceptible allele (see Methods). We found no evidence that sister species evolved resistance to fungicides (Supplementary Table S5). In the United States population pair, we found that four *CYP51* amino acids contributing to higher azole resistance increased in frequency. In Australian and Swiss population pairs, the amino acid frequencies changed towards higher resistance but the changes were not universal. Notably, at amino acid positions 188 and 513 the Swiss population pair showed changes towards the less resistant residue over time. Such changes may indicate epistasis from incompatible residues impacting protein functions (Cools et al., 2013; Hawkins et al., 2019). For resistance to MBC, we found a single amino acid residue with consistent changes towards resistance. QoI resistance emerged strongly in the Swiss population with weak or no changes in the United States and Australian pairs, respectively. We detected no shift in amino acid residues indicating resistance to SDHI as we find only a single isolate with a mutation (N86S) in the protein subunit SDH3. Overall, only the ‘hotspot’ genes for azole and MBC resistance showed at least partially correlated responses to fungicide pressure between geographical regions. Target gene mutations to QoI were not detected in Australia and SDHI resistance emerged only to a very low degree in Switzerland consistent with the historic use of each fungicide class across the continents.

### Gain in resistance generated linkage disequilibrium in resistance genes

Strong positive selection for fungicide resistance is expected to generate linkage disequilibrium in the underlying loci. To test this, we analyzed all genic and intronic SNPs in the major gene for each fungicide (Fig 3). We found strong linkage disequilibrium in *CYP51* in both the older and more recent Swiss populations consistent with the early application of azoles in Europe and the significant amino acid changes of CYP51 at the population level. The older United States population showed only a low degree of linkage disequilibrium with a substantial increase in linkage disequilibrium in the largest exon in the more recent population. The Australian population pair showed a high degree of linkage disequilibrium. Given the slow decay genome-wide, the high linkage disequilibrium in *CYP51* may stem from demographic effects rather than selection for fungicide resistance. We found relatively low degrees of linkage disequilibrium in the beta-tubulin and *SDH3* genes consistent with the lower pressure of MCB and SDHI fungicides over the analyzed time span.

**Figure 3:**
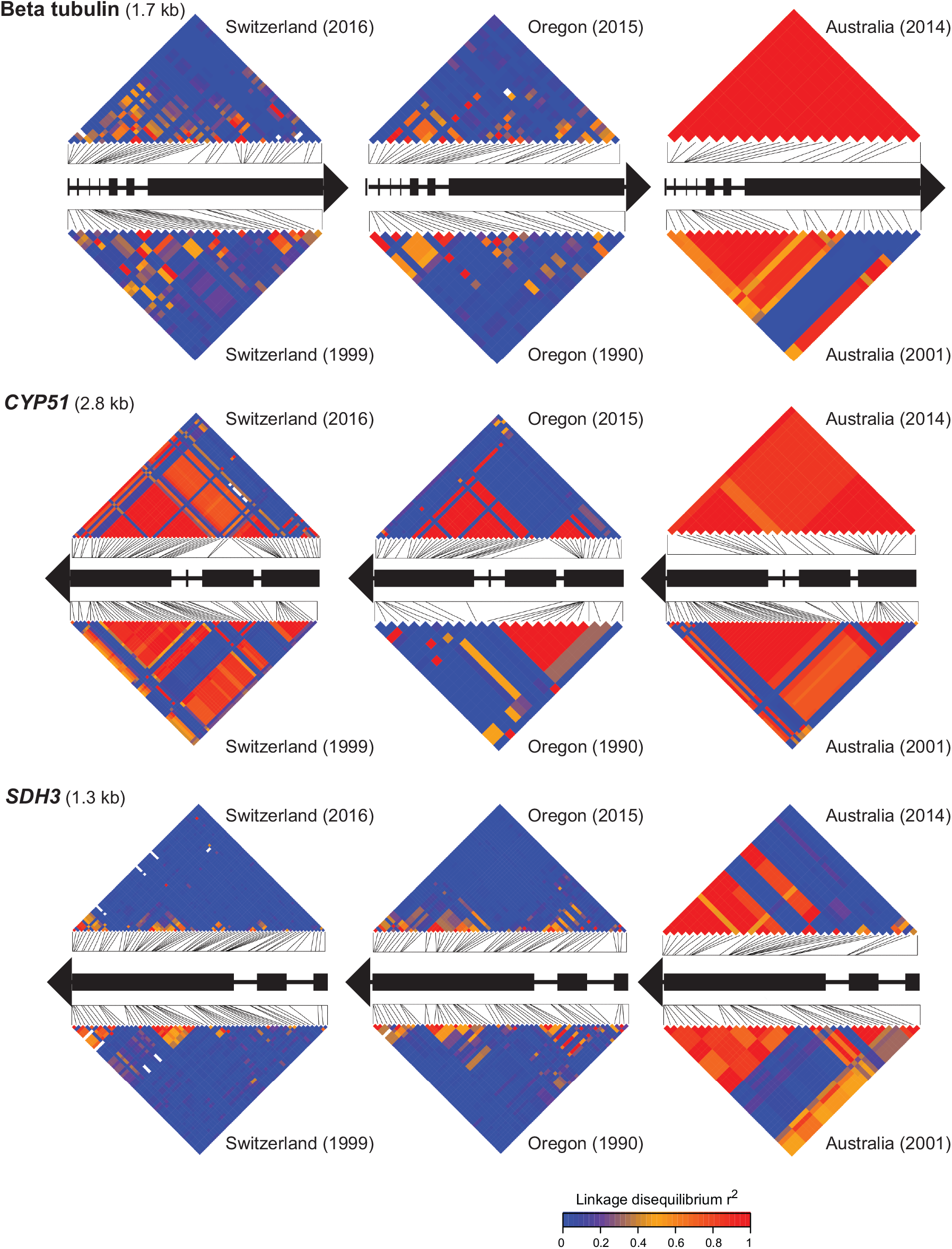
Linkage disequilibrium in the three key fungicide resistance genes. Panels show linkage disequilibrium (*r*^2^) for beta-tubulin (MCB fungicides), *CYP51* (azoles) and *SDH3* (SDHI). For each population pair, the *r*^2^ is shown for all genic and intronic SNPs both for the older (bottom heatmap) and the more recent population (top heatmap). Black rectangles represent exons.

### Excess differentiation at resistance genes among population pairs

The rise of beneficial alleles over the course of resistance evolution should lead to excess differentiation in fungicide resistance loci compared to the genomic background. We found that the average *F*_*ST*_ for resistance genes was not distinct from the genomic background in the Australian population pair (Fig 4A). However, azole resistance loci showed a substantial excess differentiation at azole resistance loci in the Swiss and US population pairs. Interestingly, *CYP51* showed the most differentiation among the Swiss population pair but the least differentiation in the US population pairs (Fig 4A). This is consistent with the finding that there were only minor amino acid frequency changes in the US population. The comparatively minor role of *CYP51* in resistance evolution in the US population pair was also shown by the GWAS where associations with resistance levels were stronger for genes other than *CYP51* (Supplementary Table S3). To account for variation in polymorphism among populations, we also estimated the absolute divergence *d*_*XY*_ (Fig 4B). The *d*_*XY*_ for *CYP51* does not show outlier values compared to the genome-wide distribution. However, the gene encoding the DHHC palmitoyl transferase shows extremely high degrees of differentiation in all three population pairs.

**Figure 4 :**
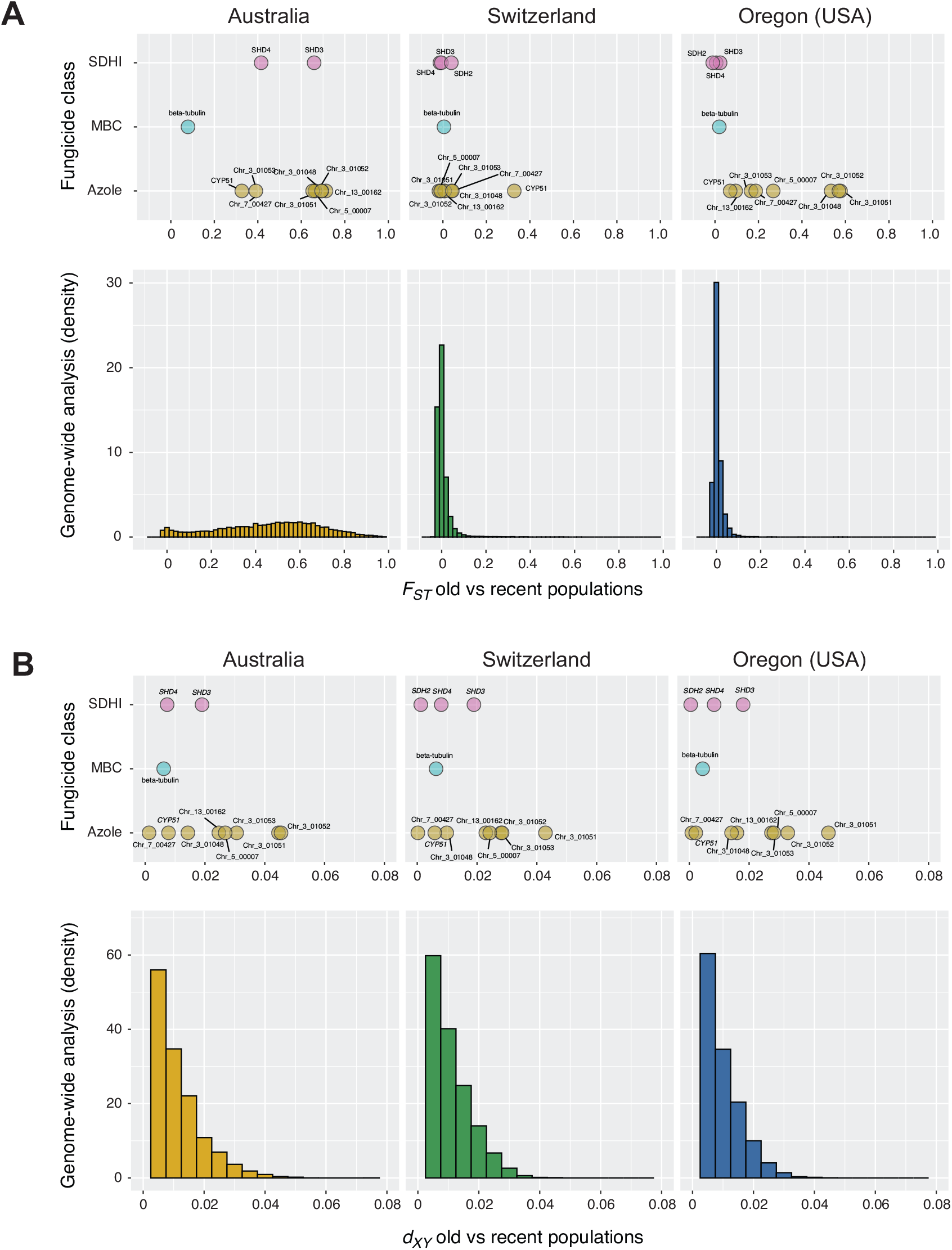
Population divergence at fungicide resistance loci in population pairs (old vs recent populations). The three population pairs in Australia, Switzerland and the United States (Oregon) are shown separately. Resistance genes are grouped according to the fungicide class and include both functionally confirmed genes (*SDH*, beta-tubulin and *CYP51*) as well as loci identified through genome-wide association studies. A) Relative divergence (*F*_*ST*_) and B) absolute divergence (*d*_*XY*_) are shown both at the level of individual resistance genes (top panel) and genome-wide (bottom panel).

### Signatures of positive selection

To directly assess whether loci underlying gains in fungicide resistance show signatures of positive selection, we performed both intra-population scans for selection as well as population pair contrasts. First, we estimated Tajima's *D* for each functionally confirmed resistance gene or genes identified by GWAS in the most recent populations. Overall, resistance loci have Tajima's *D* values as expected for the genomic background (Fig. 5A). However, most loci show a positive Tajima's *D* which suggests balancing selection or population contractions. Next, we analyzed selective sweep signatures comparing the decay of haplotypes associated with ancestral and derived alleles in the most recent populations. The iHS-based scans identified 20, 159, 93 selective sweep regions in the recent Australia, Swiss and US populations, respectively (Supplementary Fig S6A; Supplementary Table S6). Only 6.2 % of loci were shared between all three populations suggesting that recent adaptation in these populations was largely based on distinct sets of genes as previously reported (Hartmann et al., 2018). Given the genetic heterogeneity in the more recent Australian population, we repeated the selection scans excluding the eight most divergent genotypes. However, we found no impact on the number or identity of regions with signatures of selection. Next, we compared the haplotype structure within population pairs to identify the most significant changes concurrent with the gain of fungicide resistance. The XP-EHH scan identified 14, 54, 28 selective sweep regions in the Australian, Swiss and US population pairs, respectively (Fig 5B; Supplementary Fig S6B; Table S6). We found a high degree of overlap (35-42%) of selective sweep regions identified by XP-EHH and by iHS within each location. Among the functionally confirmed fungicide resistance loci, only the gene encoding beta-tubulin overlapped with a selective sweep region (Switzerland and Australia; Supplementary Table S7). In contrast, five out of seven genes identified by GWAS (excluding *CYP51*) overlapped with selective sweep regions (Switzerland and US; Supplementary Table S7). Overall, 50% of all loci associated with fungicide resistance were found in selective sweep regions in multiple populations (Fig 5C).

**Figure 5 :**
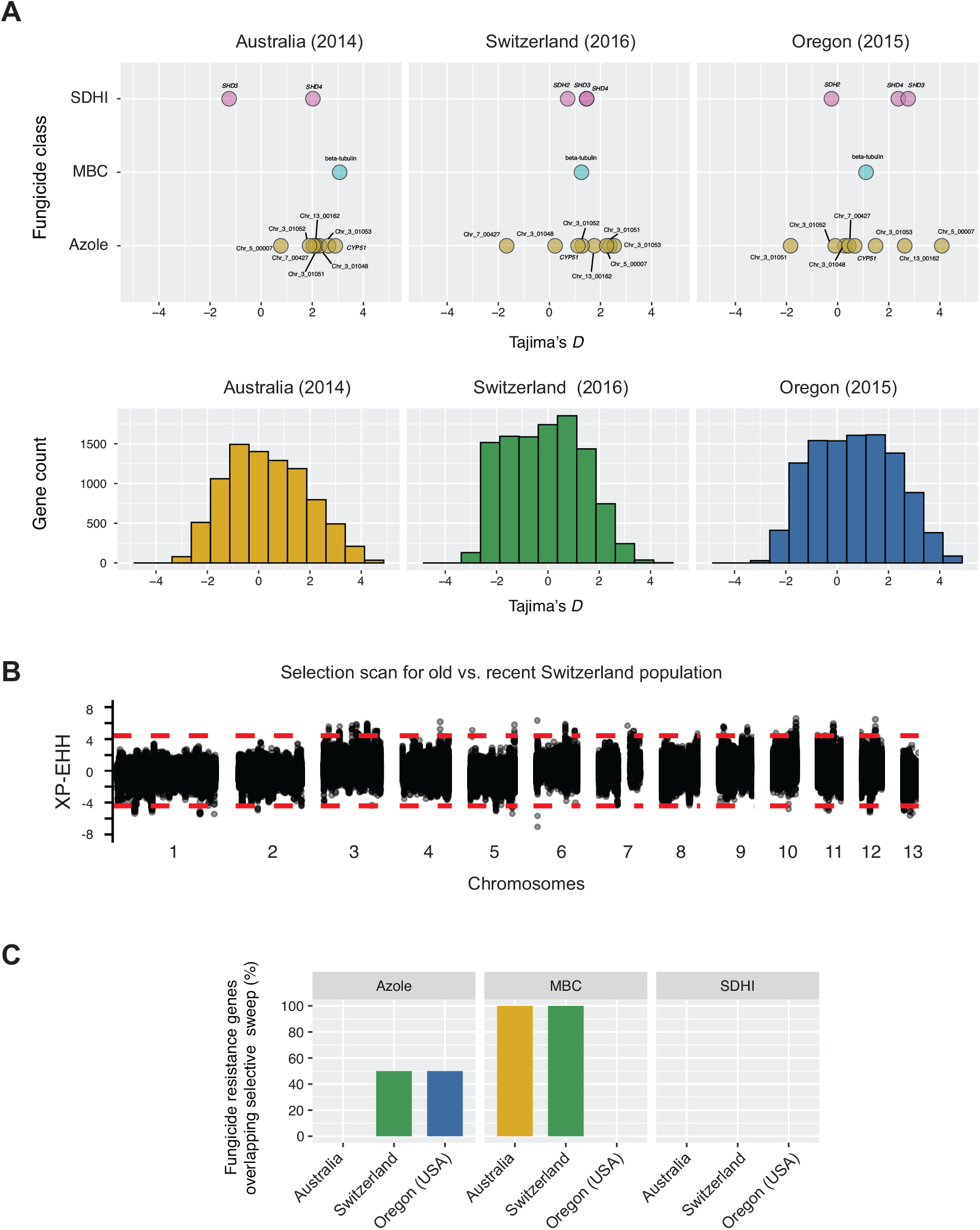
Signatures of selection in loci associated with fungicide resistance. A) Analyses of gene-wise Tajima's *D* in the recent populations. The three recent populations in Australia, Switzerland and the United States (Oregon) are shown separately. Resistance genes are grouped according to the fungicide class and include both functionally confirmed genes (*SDH*, beta-tubulin and *CYP51*) as well as loci identified through genome-wide association studies. The genome-wide distribution is shown below. B) XP-EHH scan for differences in haplotype block structure between the older and more recent populations collected in Switzerland. C) Summary of overlaps between genes associated with fungicide resistance and selective sweep regions. The percentage is shown grouped by fungicide class and geographic region.

## Discussion

Resistance to fungicides emerged rapidly and concurrently in *Z. tritici* across continents. The main drivers were “hotspots” of resistance evolution in genes encoding proteins targeted by individual fungicides. Through GWAS, we have discovered an additional major locus encoding a DHHC palmitoyl transferase with resistance gains in the US population pair and possibly supporting cell membrane stability. A large array of resistance loci showed signatures of selection, however the genes underlying the most dramatic changes largely differed among evolved populations. The application of fungicides was one of the major selective agents acting on the pathogen over the past decades.

*Z. tritici* originated in the Middle East and spread first to Europe followed by colonization events of North America and Oceania (Linde, Zhan, & McDonald, 2002; Zhan et al., 2003). We found clear population subdivisions reflecting these colonization events confirming earlier work (Hartmann et al., 2018; Zhan et al., 2003). We found that gene flow was sufficiently low among continents to produce clearly distinct genetic clusters with a high degree of homogeneity within each sampling location. The colonization of Australia was more complex with the more recent collection from Tasmania showing evidence for admixture with European genotypes influencing the trajectory of azole resistance evolution. Additionally, the Australian populations have experienced at least experienced a colonization bottleneck and possibly additional bottlenecks during the Millennium drought. Combining evidence from our own analyses of azole resistance and previous work (McDonald et al., 2019), we found that resistance increased sharply in both the United States and Australian populations. The Swiss population was already at a high resistance level at the older collection timepoint consistent with the pervasive use of azole fungicides in Europe prior to the introduction in other continents (Torriani et al., 2015). We found that the gains in resistance across continents are matched by gains in previously known resistance mutations but additional genetic factors also played a role. Using association mapping in sets of populations gaining resistance over time, we identified at least one additional major contributor to azole resistance. The DHHC palmitoyl transferase was not previously reported to confer azole resistance in plant pathogens, however analyses of *Aspergillus* fungi showed that palmitoyl transferase dysfunction can impact azole stress responses (Zhang et al., 2016). Allele frequency changes at loci identified through association mapping showed similarly drastic allele frequency shifts as loci confirmed by experimental approaches. This shows that capturing shifts in adaptive alleles requires a comprehensive to identify causal loci. In particular, geographically isolated populations may be more prone to evolve resistance based on previously unknown mutations.

A key question underlying convergent adaptation is the source of mutations (Lee & Coop, 2017) (Fig 2A). Because *Z. tritici* has colonized all major wheat-producing areas of the world prior to the application of fungicides, it is unlikely that resistance mutations originated from standing variation. Additionally, we found no evidence for resistance mutations in sister species of *Z. tritici*. Gene flow spreading resistance mutations is a likely explanation for the gain of resistance in Australia (McDonald et al., 2019). In particular, changes in amino acid residues in CYP51 associated with resistance to azoles show a striking parallelism with the European population. The parallelism extends to the two residues potentially underlying pleiotropic effects and have compensatory effects (S188N and N513K). The genome-wide analyses of neutral polymorphism support admixture as a likely source of resistance mutations in Australia. Resistance evolution in the United States population shows hallmarks of independent gains of resistance mutations with distinct mutation profiles in particular in the highly polymorphic *CYP51* gene. Strikingly, the DHHC palmitoyl transferase gene identified through GWAS showed the highest absolute divergence (*d*_*XY*_) in all three population pairs gaining azole resistance. However, *CYP51* showed the highest degree of relative divergence (*F*_*ST*_) in the Swiss population pair most likely caused by the high degree of polymorphism in the gene. The DHHC palmitoyl transferase gene showed also strong signatures of recent selection in both the United States and Switzerland populations underpinning the importance of the locus for parallel gains of fungicide resistance. Evidence for convergent adaptation across continents in *Z. tritici* establishes multiple “hotspot” genes of resistance consistent with evidence for herbicide resistance evolution (Baucom, 2019). However, the complexity of the discovered mutation shows that resistance is highly polygenic and the emergence of resistance genotypes is likely shaped by strong pleiotropy.

Convergent evolution of resistance has been a key driver of recent adaptation in *Z. tritici* and likely other plant pathogens. Polymorphism in ‘hotspot’ genes has undergone rapid allele frequency changes consistent with the heterogeneous application of different fungicide classes in both space and time. Mapping the emergence of resistance *in situ* through GWAS provided key additional loci that would have been missed by classic target gene-focused approaches. We found a higher degree of parallelism in allele frequency changes between the European and Australian populations suggesting that recent gene flow may have played a role. However, standing variation or recurrent mutations could well have been at the origin of the rise of resistance in the latest Australian sampling (McDonald et al., 2019). The United States population showed a less congruent genetic basis of resistance with the European population indicating that resistance in the United States has more likely evolved from independent mutations yet sharing most ‘hotspot’ genes. Fungicide resistance emergence shows striking parallels with the rise of herbicide resistant weeds globally. The genetic architecture of resistance in both fungi and plants shows similarly extensive variation within species largely following geographic subdivisions (Baucom, 2019). The emergence of resistance underlines the importance of understanding the ecological niche occupied by pathogens (Burdon & Thrall, 2008). Our study shows that for crop pathogens, the ability to cope with substantial input of chemicals can be a key trait for ecological success and is driven by extensive convergent evolution. Experimentally tractable microbial systems allow to simultaneously retrace the likely origin of adaptive mutations and unravel the complexity of resistance trait architectures.

## Acknowledgements

We thank Sandra Siegfried and Marcello Zala for technical assistance for DNA extraction. We are very grateful to Bruce A. McDonald, Chris Mundt and Petteri Karisto who made collections. Alice Feurtey provided helpful feedback on a previous manuscript version. AM and MCM were supported by the Australian National University, Grains and Research Development Corporation, and NSW Department of Primary Industries co-investment DAN00203 as part of the Grains, Agronomy and Pathology Partnership. This work was supported by a Marie Curie European grant (PRESTIGE-2016-4-0013) to F.E.H. F.E.H. also received the Young Biological Researcher Prize from the Fondation des Treilles, created by Anne Gruner Schlumberger, which supports research in Science and Art (https://www.les-treilles.com/la-recherche). DC was supported by the National Science Foundation (grant 31003A_173265).

## References

Arendt, J., & Reznick, D. (2008). Convergence and parallelism reconsidered: what have we learned about the genetics of adaptation? Trends in Ecology & Evolution, 23(1), 26–32.

Azevedo, M.-M., Faria-Ramos, I., Cruz, L. C., Pina-Vaz, C., & Rodrigues, A. G. (2015). Genesis of Azole Antifungal Resistance from Agriculture to Clinical Settings. Journal of Agricultural and Food Chemistry, 63(34), 7463–7468.

Baucom, R. S. (2019). Evolutionary and ecological insights from herbicide-resistant weeds: what have we learned about plant adaptation, and what is left to uncover? The New Phytologist, 223(1), 68–82.

Bolger, A. M., Lohse, M., & Usadel, B. (2014). Trimmomatic: a flexible trimmer for Illumina sequence data. Bioinformatics, 30(15), 2114–2120.

Bolnick, D. I., Barrett, R. D. H., Oke, K. B., Rennison, D. J., & Stuart, Y. E. (2018). (Non)Parallel Evolution. Annual Review of Ecology, Evolution, and Systematics, Vol. 49, pp. 303–330. doi: 10.1146/annurev-ecolsys-110617-062240

Bradbury, P. J., Zhang, Z., Kroon, D. E., Casstevens, T. M., Ramdoss, Y., & Buckler, E. S. (2007). TASSEL: software for association mapping of complex traits in diverse samples. Bioinformatics, 23(19), 2633–2635.

Burdon, J. J., & Thrall, P. H. (2008). Pathogen evolution across the agro-ecological interface: implications for disease management. Evolutionary Applications, 1(1), 57–65.

Chan, Y. F., Marks, M. E., Jones, F. C., Villarreal, G., Jr, Shapiro, M. D., Brady, S. D., … Kingsley, D. M. (2010). Adaptive evolution of pelvic reduction in sticklebacks by recurrent deletion of a Pitx1 enhancer. Science, 327(5963), 302–305.

Colosimo, P. F. (2005). Widespread Parallel Evolution in Sticklebacks by Repeated Fixation of Ectodysplasin Alleles. Science, Vol. 307, pp. 1928–1933. doi: 10.1126/science.1107239

Cools, H. J., Bayon, C., Atkins, S., Lucas, J. A., & Fraaije, B. A. (2012). Overexpression of the sterol 14α-demethylase gene (MgCYP51) in Mycosphaerella graminicola isolates confers a novel azole fungicide sensitivity phenotype. Pest Management Science, 68(7), 1034–1040.

Cools, H. J., & Fraaije, B. A. (2013). Update on mechanisms of azole resistance in Mycosphaerella graminicola and implications for future control. Pest Management Science, 69(2), 150–155.

Cools, H. J., Hawkins, N. J., & Fraaije, B. A. (2013). Constraints on the evolution of azole resistance in plant pathogenic fungi. Plant Pathology, Vol. 62, pp. 36–42. doi: 10.1111/ppa.12128

Cools, H. J., Mullins, J. G. L., Fraaije, B. A., Parker, J. E., Kelly, D. E., Lucas, J. A., & Kelly, S. L. (2011). Impact of recently emerged sterol 14{alpha}-demethylase (CYP51) variants of Mycosphaerella graminicola on azole fungicide sensitivity. Applied and Environmental Microbiology, 77(11), 3830–3837.

Cowger, C., Hoffer, M. E., & Mundt, C. C. (2000). Specific adaptation by Mycosphaerella graminicola to a resistant wheat cultivar. Plant Pathology, Vol. 49, pp. 445–451. doi: 10.1046/j.1365-3059.2000.00472.x

Délye, C., Jasieniuk, M., & Le Corre, V. (2013). Deciphering the evolution of herbicide resistance in weeds. Trends in Genetics: TIG, 29(11), 649–658.

Estep, L. K., Torriani, S. F. F., Zala, M., Anderson, N. P., Flowers, M. D., McDonald, B. A., … Brunner, P. C. (2015). Emergence and early evolution of fungicide resistance in North American populations ofZymoseptoria tritici. Plant Pathology, Vol. 64, pp. 961–971. doi: 10.1111/ppa.12314

Fisher, M. C., Henk, D. A., Briggs, C. J., Brownstein, J. S., Madoff, L. C., McCraw, S. L., & Gurr, S. J. (2012). Emerging fungal threats to animal, plant and ecosystem health. Nature, 484(7393), 186–194.

Fukata, M., Fukata, Y., Adesnik, H., Nicoll, R. A., & Bredt, D. S. (2004). Identification of PSD-95 palmitoylating enzymes. Neuron, 44(6), 987–996.

Gautier, M., Klassmann, A., & Vitalis, R. (2017). rehh2.0: a reimplementation of the R packagerehhto detect positive selection from haplotype structure. Molecular Ecology Resources, Vol. 17, pp. 78–90. doi: 10.1111/1755-0998.12634

Goodwin, S. B., M’barek, S. B., Dhillon, B., Wittenberg, A. H. J., Crane, C. F., Hane, J. K., … Kema, G. (2011). Finished genome of the fungal wheat pathogen Mycosphaerella graminicola reveals dispensome structure, chromosome plasticity, and stealth pathogenesis. PLoS Genetics, 7(6), e1002070.

Hartmann, F. E., McDonald, B. A., & Croll, D. (2018). Genome-wide evidence for divergent selection between populations of a major agricultural pathogen. Molecular Ecology, 27(12), 2725–2741.

Hartmann, F. E., Sánchez-Vallet, A., McDonald, B. A., & Croll, D. (2017). A fungal wheat pathogen evolved host specialization by extensive chromosomal rearrangements. The ISME Journal, 11(5), 1189–1204.

Hawkins, N. J., Bass, C., Dixon, A., & Neve, P. (2019). The evolutionary origins of pesticide resistance. Biological Reviews, Vol. 94, pp. 135–155. doi: 10.1111/brv.12440

Heliconius Genome Consortium. (2012). Butterfly genome reveals promiscuous exchange of mimicry adaptations among species. Nature, 487(7405), 94–98.

Jakobsson, M., & Rosenberg, N. A. (2007). CLUMPP: a cluster matching and permutation program for dealing with label switching and multimodality in analysis of population structure. Bioinformatics, 23(14), 1801–1806.

Langmead, B., & Salzberg, S. L. (2012). Fast gapped-read alignment with Bowtie 2. Nature Methods, 9(4), 357–359.

Lee, K. M., & Coop, G. (2017). Distinguishing Among Modes of Convergent Adaptation Using Population Genomic Data. Genetics, 207(4), 1591–1619.

Linde, C. C., Zhan, J., & McDonald, B. A. (2002). Population Structure of Mycosphaerella graminicola: From Lesions to Continents. Phytopathology, 92(9), 946–955.

Linder, M. E., & Deschenes, R. J. (2007). Palmitoylation: policing protein stability and traffic. Nature Reviews. Molecular Cell Biology, 8(1), 74–84.

Losos, J. B. (2011). CONVERGENCE, ADAPTATION, AND CONSTRAINT. Evolution, Vol. 65, pp. 1827–1840. doi: 10.1111/j.1558-5646.2011.01289.x

Lucas, J. A., Hawkins, N. J., & Fraaije, B. A. (2015). The Evolution of Fungicide Resistance. Advances in Applied Microbiology, pp. 29–92. doi: 10.1016/bs.aambs.2014.09.001

Martin, A., & Orgogozo, V. (2013). The Loci of repeated evolution: a catalog of genetic hotspots of phenotypic variation. Evolution; International Journal of Organic Evolution, 67(5), 1235–1250.

McDonald, M. C., Renkin, M., Spackman, M., Orchard, B., Croll, D., Solomon, P. S., & Milgate, A. (2019). Rapid Parallel Evolution of Azole Fungicide Resistance in Australian Populations of the Wheat Pathogen. Applied and Environmental Microbiology, 85(4). doi: 10.1128/AEM.01908-18

McKenna, A., Hanna, M., Banks, E., Sivachenko, A., Cibulskis, K., Kernytsky, A., … DePristo, M. A. (2010). The Genome Analysis Toolkit: a MapReduce framework for analyzing next-generation DNA sequencing data. Genome Research, 20(9), 1297–1303.

Meile, L., Croll, D., Brunner, P. C., Plissonneau, C., Hartmann, F. E., McDonald, B. A., & Sánchez-Vallet, A. (2018). A fungal avirulence factor encoded in a highly plastic genomic region triggers partial resistance to septoria tritici blotch. New Phytologist, Vol. 219, pp. 1048–1061. doi: 10.1111/nph.15180

Menchari, Y., Camilleri, C., Michel, S., Brunel, D., Dessaint, F., Le Corre, V., & Délye, C. (2006). Weed response to herbicides: regional-scale distribution of herbicide resistance alleles in the grass weed Alopecurus myosuroides. The New Phytologist, 171(4), 861–873.

Mohd-Assaad, N., McDonald, B. A., & Croll, D. (2016). Multilocus resistance evolution to azole fungicides in fungal plant pathogen populations. Molecular Ecology, Vol. 25, pp. 6124–6142. doi: 10.1111/mec.13916

Molin, W. T., Yaguchi, A., Blenner, M. A., & Saski, C. A. (2020). The eccDNA Replicon: A Heritable, Extra-Nuclear Vehicle that Enables Gene Amplification and Glyphosate Resistance in Amaranthus palmeri. The Plant Cell. doi: 10.1105/tpc.20.00099

Oggenfuss, U., Badet, T., Wicker, T., & Hartmann, F. E. (2020). A population-level invasion by transposable elements in a fungal pathogen. BioRxiv. Retrieved from https://www.biorxiv.org/content/10.1101/2020.02.11.944652v1.abstract

Omrane, S., Audéon, C., Ignace, A., Duplaix, C., Aouini, L., Kema, G., … Fillinger, S. (2017). Plasticity of the Promoter Leads to Multidrug Resistance in the Wheat Pathogen. mSphere, 2(5). doi: 10.1128/mSphere.00393-17

Omrane, S., Sghyer, H., Audéon, C., Lanen, C., Duplaix, C., Walker, A.-S., & Fillinger, S. (2015). Fungicide efflux and the MgMFS1 transporter contribute to the multidrug resistance phenotype in Zymoseptoria tritici field isolates. Environmental Microbiology, 17(8), 2805–2823.

Orr, H. A., & Allen Orr, H. (2005).The probability of parallel evolution. Evolution, Vol. 59, pp. 216–220. doi: 10.1111/j.0014-3820.2005.tb00907.x

Parker, J. E., Warrilow, A. G. S., Price, C. L., Mullins, J. G. L., Kelly, D. E., & Kelly, S. L. (2014). Resistance to antifungals that target CYP51. Journal of Chemical Biology, 7(4), 143–161.

Pearce, R. J., Pota, H., Evehe, M.-S. B., Bâ, E.-H., Mombo-Ngoma, G., Malisa, A. L., … Roper, C. (2009). Multiple origins and regional dispersal of resistant dhps in African Plasmodium falciparum malaria. PLoS Medicine, 6(4), e1000055.

Pfeifer, B., Wittelsbürger, U., Ramos-Onsins, S. E., & Lercher, M. J. (2014). PopGenome: an efficient Swiss army knife for population genomic analyses in R. Molecular Biology and Evolution, 31(7), 1929–1936.

Powles, S. B., & Yu, Q. (2010). Evolution in action: plants resistant to herbicides. Annual Review of Plant Biology, 61, 317–347.

Purcell, S., Neale, B., Todd-Brown, K., Thomas, L., Ferreira, M. A. R., Bender, D., … Sham, P. C. (2007). PLINK: a tool set for whole-genome association and population-based linkage analyses. American Journal of Human Genetics, 81(3), 559–575.

Raj, A., Stephens, M., & Pritchard, J. K. (2014). fastSTRUCTURE: variational inference of population structure in large SNP data sets. Genetics, 197(2), 573–589.

Rehfus, A., Strobel, D., Bryson, R., & Stammler, G. (2018). Mutations in sdh genes in field isolates of Zymoseptoria tritici and impact on the sensitivity to various succinate dehydrogenase inhibitors. Plant Pathology, 67(1), 175–180.

Roesti, M., Gavrilets, S., Hendry, A. P., Salzburger, W., & Berner, D. (2014). The genomic signature of parallel adaptation from shared genetic variation. Molecular Ecology, 23(16), 3944–3956.

Sabeti, P. C., Varilly, P., Fry, B., Lohmueller, J., Hostetter, E., Cotsapas, C., … Stewart, J. (2007). Genome-wide detection and characterization of positive selection in human populations. Nature, 449(7164), 913–918.

Shimizu, M., Goto, M., Hanai, M., Shimizu, T., Izawa, N., Kanamoto, H., … Kobayashi, H. (2008). Selectable tolerance to herbicides by mutated acetolactate synthase genes integrated into the chloroplast genome of tobacco. Plant Physiology, 147(4), 1976–1983.

Singh, N. K., Chanclud, E., & Croll, D. (2020). Population-level deep sequencing reveals the interplay of clonal and sexual reproduction in the fungal wheat pathogen Zymoseptoria tritici. doi: 10.1101/2020.07.07.191510

Song, Y., Endepols, S., Klemann, N., Richter, D., Matuschka, F.-R., Shih, C.-H., … Kohn, M. H. (2011). Adaptive Introgression of Anticoagulant Rodent Poison Resistance by Hybridization between Old World Mice. Current Biology, Vol. 21, pp. 1296–1301. doi: 10.1016/j.cub.2011.06.043

Stern, D. L. (2013). The genetic causes of convergent evolution. Nature Reviews. Genetics, 14(11), 751–764.

Storey, J. D., Bass, A. J., Dabney, A., & Robinson, D. (2015). qvalue: Q-value estimation for false discovery rate control. R package version 2.0. 0. Available at Github.Com/jdstorey/qvalue. Accessed April, 14, 2017.

Stukenbrock, E. H., Bataillon, T., Dutheil, J. Y., Hansen, T. T., Li, R., Zala, M., … Schierup, M. H. (2011). The making of a new pathogen: insights from comparative population genomics of the domesticated wheat pathogen Mycosphaerella graminicola and its wild sister species. Genome Research, 21(12), 2157–2166.

Stukenbrock, E. H., Jørgensen, F. G., Zala, M., Hansen, T. T., McDonald, B. A., & Schierup, M. H. (2010). Whole-genome and chromosome evolution associated with host adaptation and speciation of the wheat pathogen Mycosphaerella graminicola. PLoS Genetics, 6(12), e1001189.

Tishkoff, S. A., Reed, F. A., Ranciaro, A., Voight, B. F., Babbitt, C. C., Silverman, J. S., … Deloukas, P. (2007). Convergent adaptation of human lactase persistence in Africa and Europe. Nature Genetics, 39(1), 31–40.

Torriani, S., Brunner, P. C., McDonald, B. A., & Sierotzki, H. (2009). QoI resistance emerged independently at least 4 times in European populations of Mycosphaerella graminicola. Pest Management Science, 65(2), 155–162.

Torriani, S. F. F., Melichar, J. P. E., Mills, C., Pain, N., Sierotzki, H., & Courbot, M. (2015). Zymoseptoria tritici: A major threat to wheat production, integrated approaches to control. Fungal Genetics and Biology: FG & B, 79, 8–12.

Tranel, P. J., Wright, T. R., & Heap, I. M. (n.d.). Mutations in herbicide-resistant weeds to ALS inhibitors. International Survey of Herbicide Resistant Weeds. Retrieved July 16, 2020, from www.weedscience.org

Zhang, Y., Zheng, Q., Sun, C., Song, J., Gao, L., Zhang, S., … Lu, L. (2016). Palmitoylation of the Cysteine Residue in the DHHC Motif of a Palmitoyl Transferase Mediates Ca2+ Homeostasis in Aspergillus. PLoS Genetics, 12(4), e1005977.

Zhan, J., Pettway, R. E., & McDonald, B. A. (2003). The global genetic structure of the wheat pathogen Mycosphaerella graminicola is characterized by high nuclear diversity, low mitochondrial diversity, regular recombination, and gene flow. Fungal Genetics and Biology: FG & B, 38(3), 286–297.

Zhong, Z., Marcel, T. C., Hartmann, F. E., Ma, X., Plissonneau, C., Zala, M., … Palma-Guerrero, J. (2017). A small secreted protein in Zymoseptoria tritici is responsible for avirulence on wheat cultivars carrying the Stb6 resistance gene. The New Phytologist, 214(2), 619–631.

