## Supplementary Figure for "The complex genomic basis of rapid convergent adaptation to pesticides across continents in a fungal plant pathogen"

#### Supplementary Tables

(see separate Excel file combining all tables)

**Supplementary Table S1 : List of the 356 analyzed *Zymoseptoria tritici* isolates including location and year of sampling and NCBI SRA BioProject accession numbers of genome data.**

**Supplementary Table S2 : Non-synonymous mutations in six resistance genes reported to confer increased fungicide resistance.**

**Supplementary Table S3 : List of significant SNPs identified by GWAS for resistance to propiconazole**. A. GWAS performed on a set of 211 isolates from the United States (Oregon), Australia, Switzerland and Israel. B. GWAS performed on a set of 134 isolates from the United States (Oregon) including the older and more recent collection (1990 and 2015). Gene *Zt09_chr_3_01051*, which encodes a DHHC palmitoyl transferase is highlighted in blue.

**Supplementary Table S4 : Functional description of genes encoded in the eight significantly association GWAS loci for propiconazole resistance.** Pangenome characteristics and functional descriptions were retrieved from Badet et al. (2020).

**Supplementary Table S5 : Fungicide resistance genotypes in the 356 *Zymoseptoria tritici* isolates across major resistance loci.** Non-synonymous substitutions are shown grouped by gene.

**Supplementary Table S6 : Description of selective sweep regions identified by iHS scans in the more recent populations collected in Australia, Switzerland and the United States (Oregon).**

**Supplementary Table S7 : Description of selective sweep regions identified by XP-EHH scans contrasting haplotype structures between pairs of older and more recent populations collected in Switzerland, Australia and the United States (Oregon).**

**Supplementary Table S8 : Overview of loci linked to fungicide resistance and found in a selective sweep region.**

### Supplementary Figures

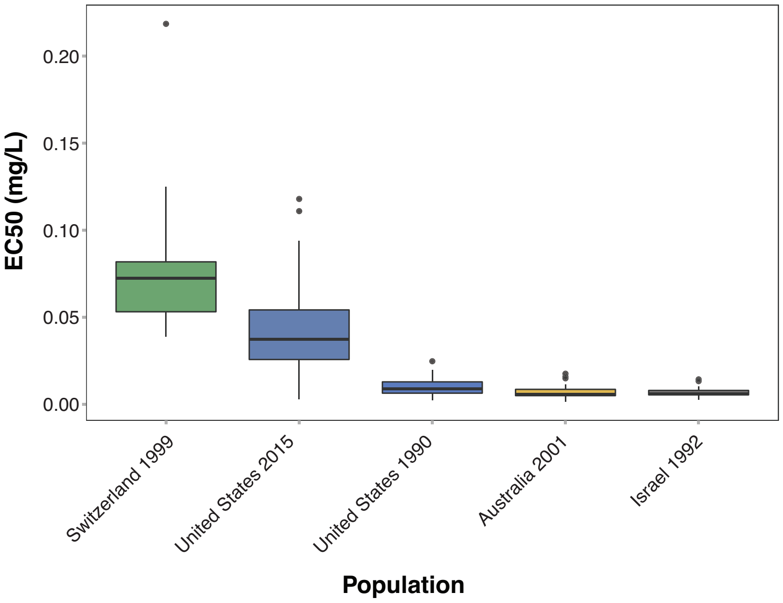


**Supplementary Figure S1 : Variation in propiconazole resistance among populations.** Distribution of half-maximum concentrations (EC50) for isolates from the older and more recent United States populations, as well as the older populations from Switzerland, Israel and Australia.

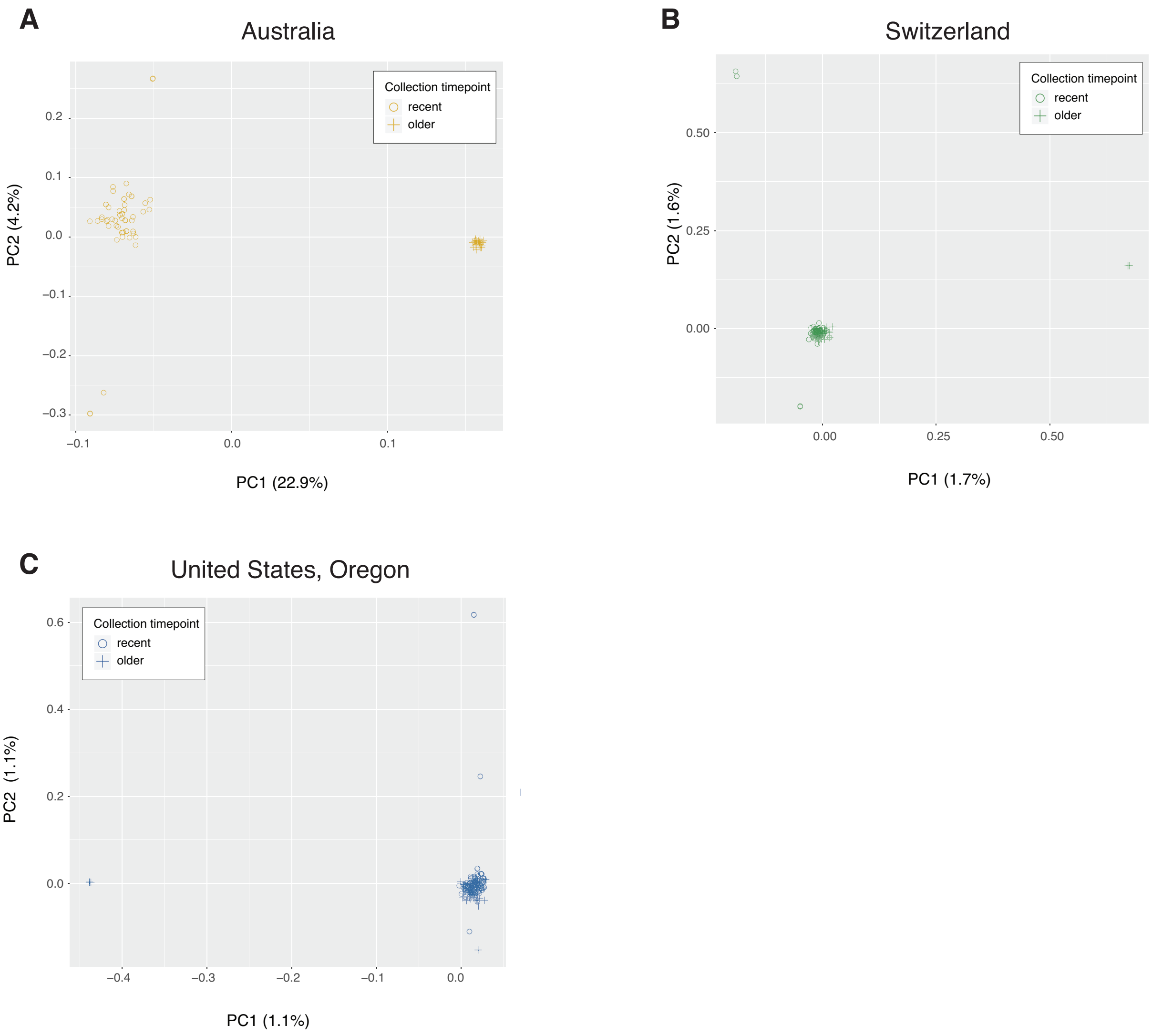


**Supplementary Figure S2 : Fine-scale population structure within geographical region.** Principal component analyses (PCA) for isolates sampled in A) Australia, B) Switzerland, and C) the United States (Oregon).

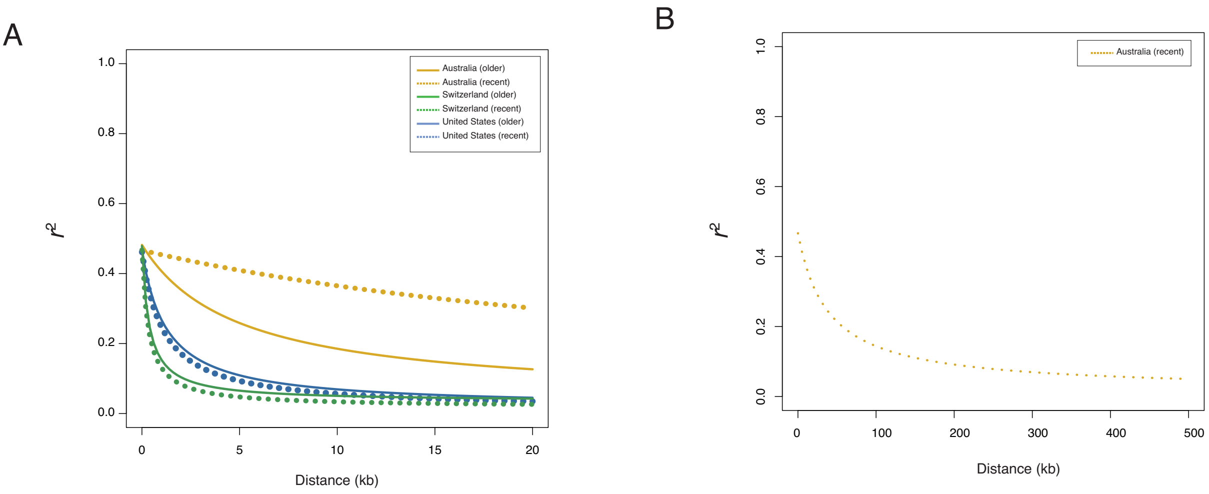


**Supplementary Figure S3 : Linkage disequilibrium decay in populations.** A) Decay of $r^2$ according to genomic distance in the six populations for all pairs of SNP loci up to a maximum distance of 20 kb. B) Decay of $r^2$ according to genomic distance in the more recent population from Australia for pairs of SNPs sampled at a maximum distance of 500 kb.

A

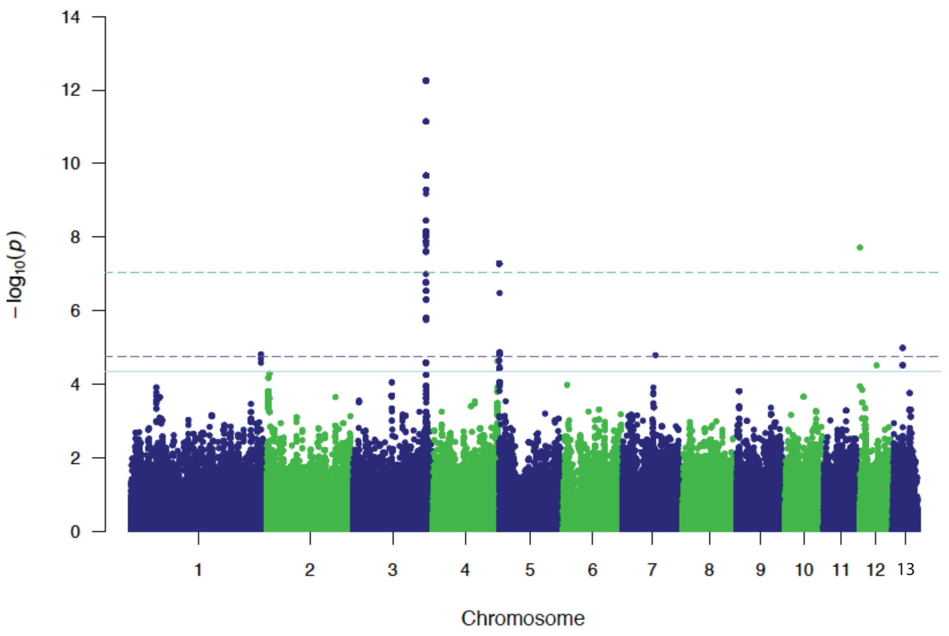


B

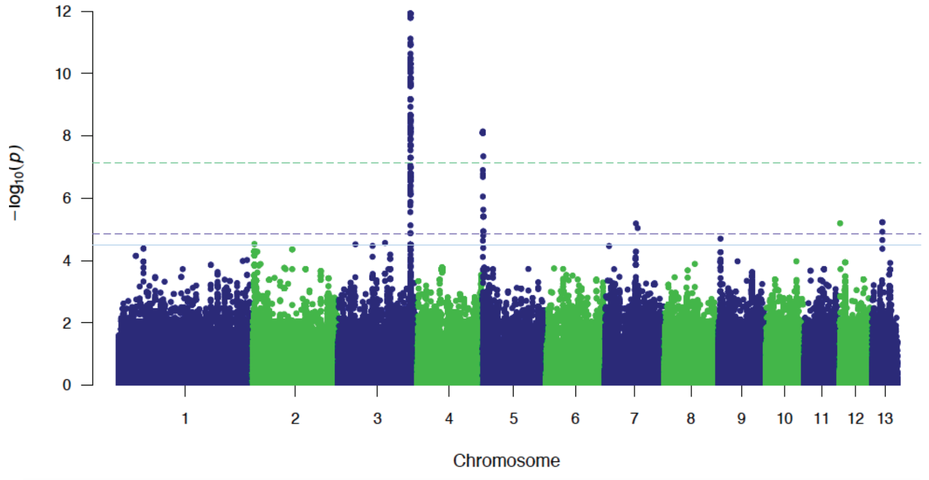


**Supplementary Figure S4 : Genome-wide association mapping for resistance to propiconazoles**. A) Manhattan plot of the GWAS analyses for the combined older and more recent populations from the United States (Oregon; $n$ = 134). B) Manhattan plot of the GWAS analyses including a larger set of isolates from the older and more recent United States populations, as well as the older populations from Switzerland, Israel and Australia ($n$ = 211). The Bonferroni (alpha = 0.05), FDR 5% and FDR 10% thresholds are shown with dotted green, dotted blue and solid blue lines, respectively.

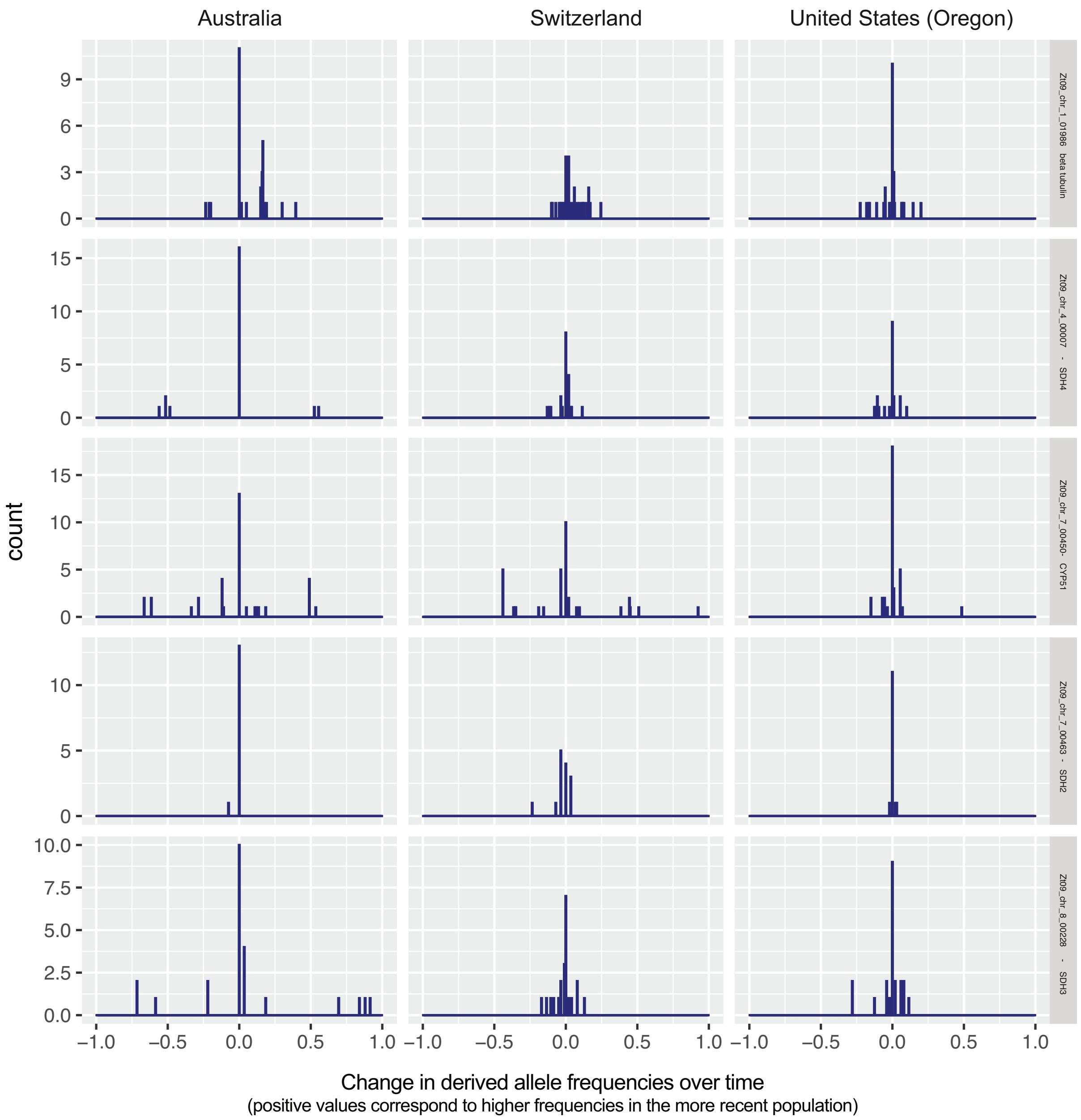


**Supplementary Figure S5: Distribution of derived allele frequency changes for SNPs contained in genes linked to fungicide resistance.** Loci were sorted out by fungicide class: azoles, MCB and SDHIs. For GWAS candidate loci on chromosome 3, only gene *Zt09_chr_3_01051* predicted to encode a DHHC palmitoyl transferase was shown. **5A): Parallel changes in allele frequencies in response to fungicide pressure**. SNPs in functionally confirmed fungicide resistance genes. Allele frequency changes are expressed as the difference in allele frequencies between older and more recent populations. Positive values correspond to higher frequencies of the derived allele in the more recent populations.

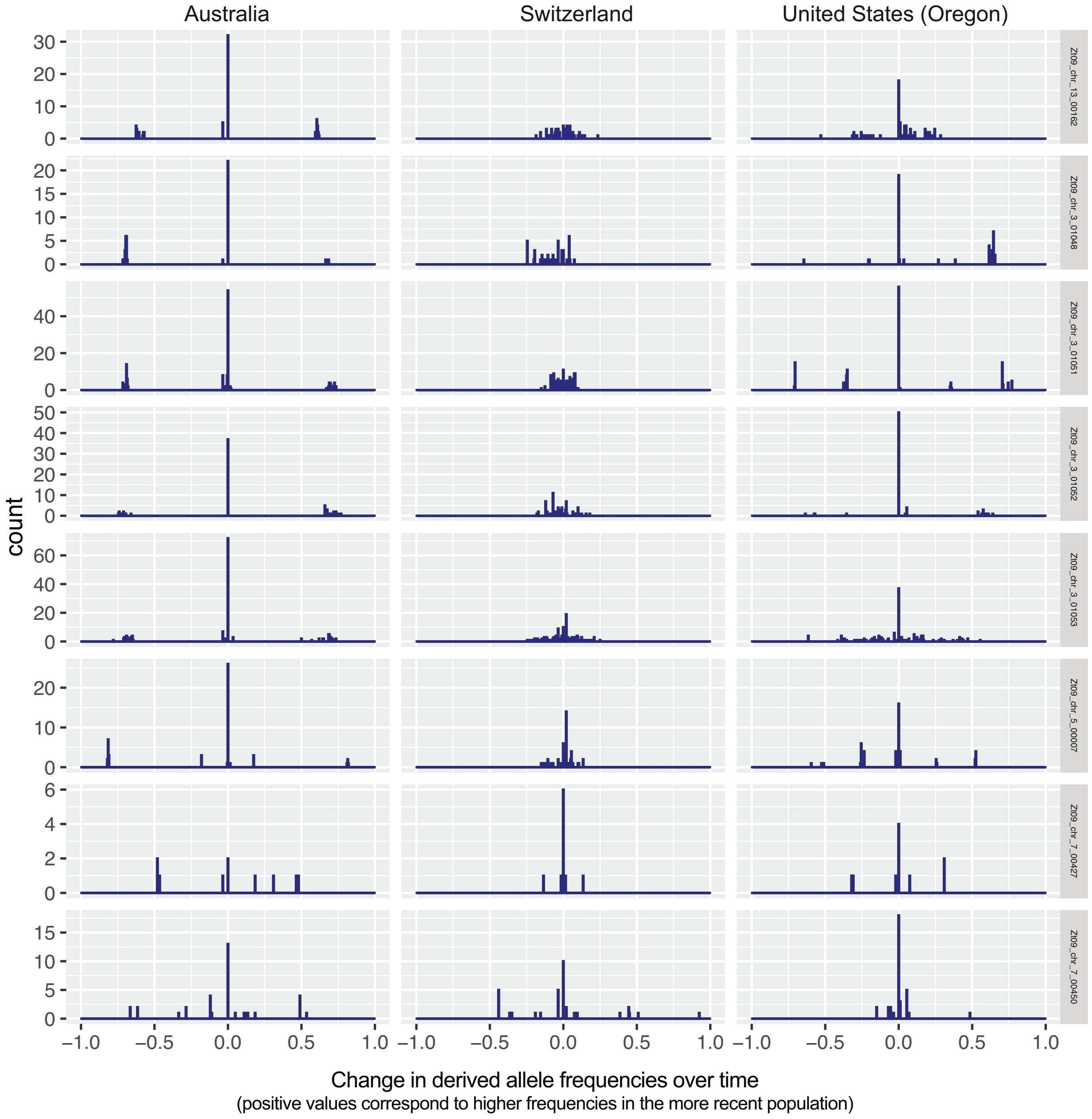


**Supplementary Figure 5B: Parallel changes in allele frequencies in response to fungicide pressure**. SNPs in fungicide resistance genes identified through genome-wide association mapping. Allele frequency changes are expressed as the difference in allele frequencies between older and more recent populations. Positive values correspond to higher frequencies of the derived allele in the more recent populations.

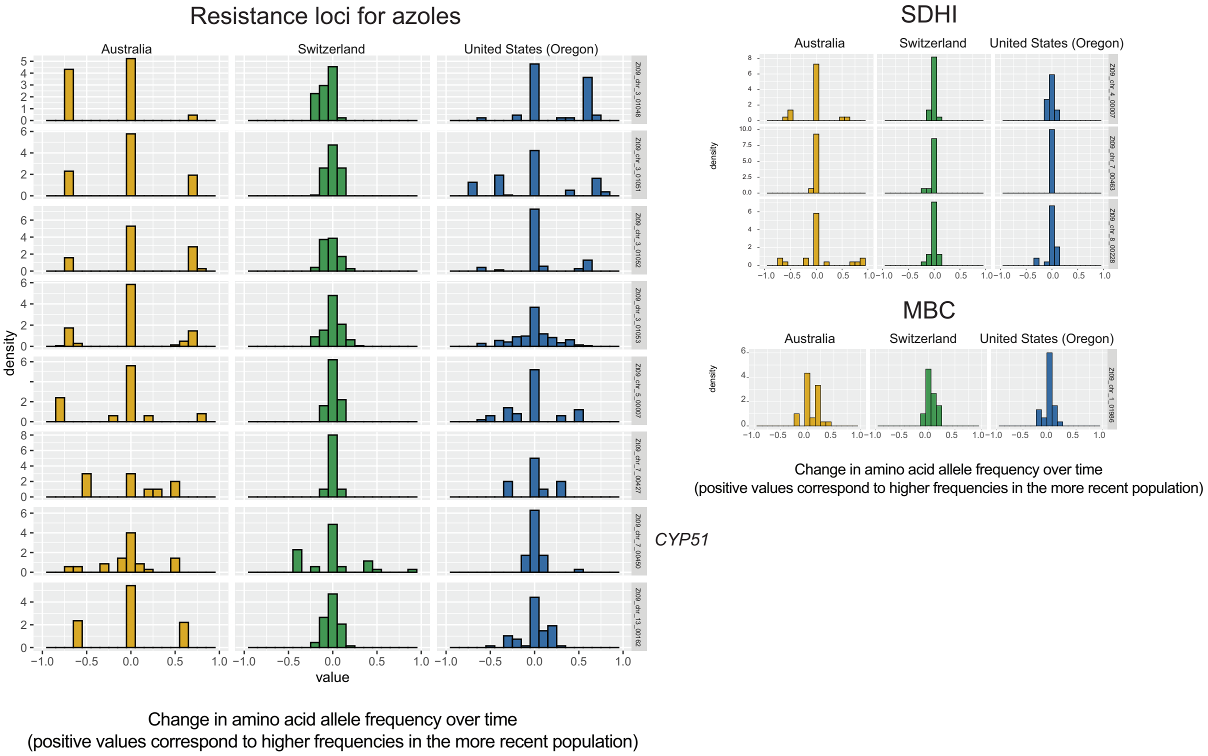


**Supplementary Figure 5C: Parallel changes in allele frequencies in response to fungicide pressure**. Changes in amino acid frequencies in populations between older and more recent populations. Positive frequency changes correspond to gains in the amino acid associated with higher resistance in the more recent populations.

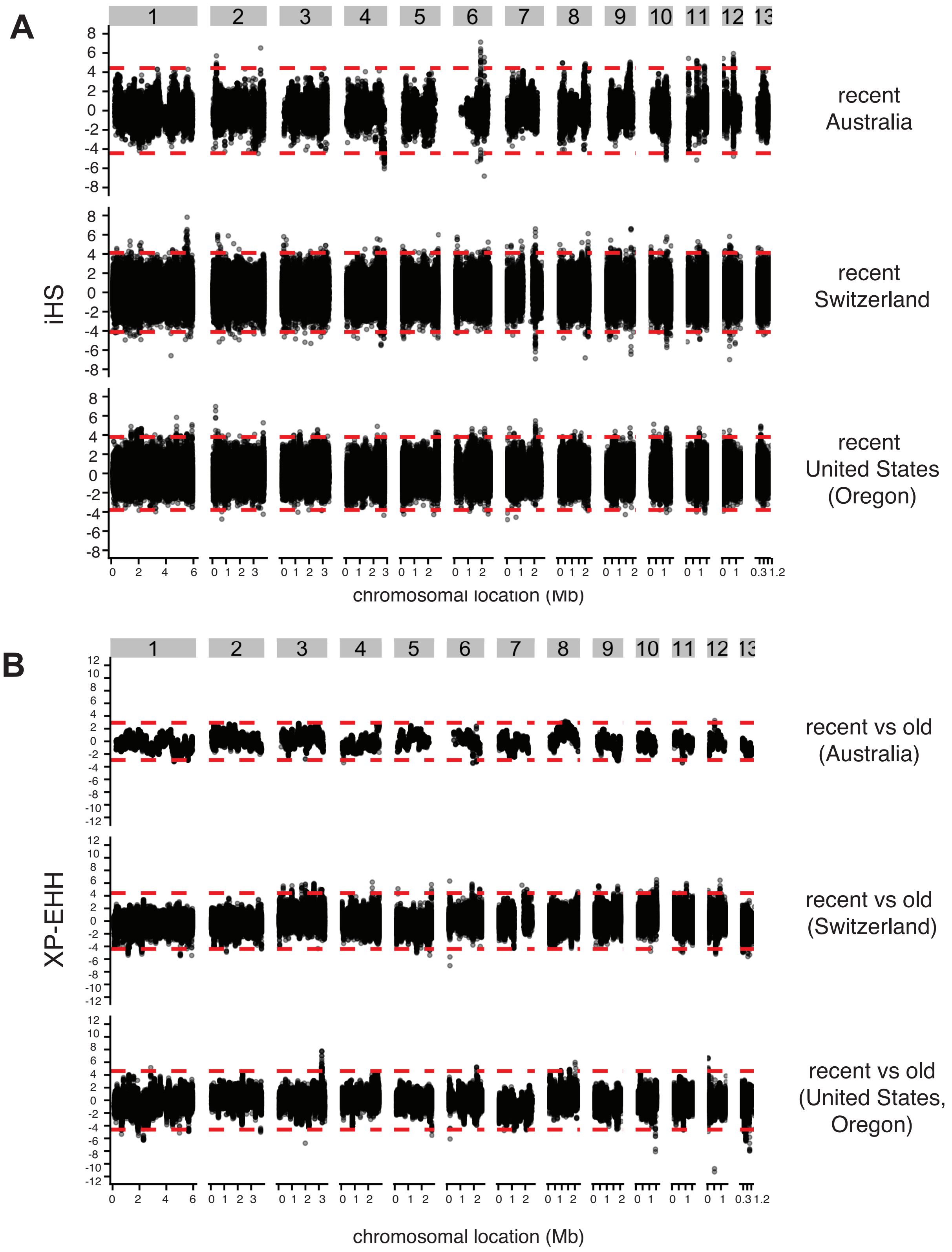


**Supplementary Figure S6 : Selective sweeps identified through haplotype structure analyses.** A) iHS scans performed separately for each of the more recent populations. Dotted horizontal lines correspond to the lower and upper 99.9 percentiles of the iHS statistic. B) XP-EHH scans performed between recent and old populations separately for each of the three geographical regions. Dotted horizontal lines correspond to the lower and upper 99.9 percentiles of the XP-EHH statistic.
